# Biomechanical Analysis of Angular Motion in Association with Bilateral Semicircular Canal Function

**DOI:** 10.1101/677872

**Authors:** Shuang Shen, Fei Zhao, Zhaoyue Chen, Qingyin Zheng, Shen Yu, Tongtao Cao, Peng Ma

## Abstract

The aim of this study was to develop a finite element (FE) model of bilateral human semicircular canals (SCCs) in order to simulate and analyze the complex fluid-structural interaction between the endolymph and cupulae by calculating the degree of cupular expansion and the cupular deflection. The results showed that cupular deflection responses were consistent with Ewald’s II law, whereas each pair of bilateral cupulae simultaneously expanded or compressed to the same degree. In addition, both the degree of cupular expansion and cupular deflection can be expressed as the solution of forced oscillation during head sinusoidal rotation, and the amplitude of cupular expansion was approximately two times greater than that of cupular deflection. Regarding the amplitude-frequency and phase-frequency characteristics, the amplitude ratios among the horizontal semicircular canal (HC) cupular expansion, the anterior semicircular canal (AC) cupular expansion, and the posterior semicircular canal (PC) cupular expansion was constant at 1:0.82:1.62, and the phase differences among them were constant at 0 or 180 degrees at the frequencies of 0.5 to 6 Hz. However, both the amplitude ratio and the phase differencies of the cupular deflection incresed nonlinearly with the increase of frequency and tended to be constant at the frequency band between 2 and 6 Hz. The results indicate that the responses of cupular expansion might only be related to the mass and rigidity of three cupulae and the endolymph, but the responses of cupular deflection are related to the mass, rigidity, or damping of them, and these physical properties would be affected by vestibular dysfunction. Therefore, both the degree of cupular expansion and cupular deflection should be considered important mechanical variables for induced neural signals. Such a numerical model can be further built to provide a useful theoretical approach for exploring the biomechanical nature underlying vestibular dysfunction.

**Statement of significance:** By taking the advantage of the torsional pendulum model and the FE model, a healthy human vestibular SCCs was developed to investigate the angular motion in association with SCC function. As a result, the responses of cupular expansion and deflection during head horizontal sinusoidal rotation were analyzed for the first time, showing quantitative correlation to the eye movement due to the vestibular ocular reflex (VOR) pathway. These responses play important roles in the cupular mechano-electrical transduction process. The significant outcome derived from this study provides a useful theoretical approach for further exploring the biomechanical nature underlying vestibular dysfunction.

## Introduction

The peripheral vestibular system is part of the inner ear, including three pairs of semicircular canals (SCCs) together with the utricle and saccule. It plays a primary role in the maintenance of both postural stability and stabilization of the visual image on the retina. The mechanism underlying the vestibular system function is generally considered a three-dimensional inertial-guidance system, in which the SCCs are known to detect any rotational motion of the head, while the utricle and saccule perceive linear acceleration and gravity. Three pairs of SCCs are approximately symmetrical to each other along the central sagittal plane, with three ipsilateral SCCs mutually orthogonal to each other. The sensory epithelium resides in the ampulla of the membranous SCCs, which are a toroidal loop of the endolymph and eventually assemble in the utricle. When the head undergoes rotation, the inertia simultaneously forces endolymphatic flow and cupular deformation, and then cupulae drag the stereocilia and kinocilia bend. As a result, intricate mechano-electrical transduction induces neural signals to enable the brain to determine the magnitude and direction of the head rotation and to stabilize the gaze through the vestibular ocular reflex (VOR) (1).

Currently, there is a series of clinical tests for examining the vestibular system function, such as the caloric test (2), the rotational chair test (RCT) (3), and the vestibular autorotation test (VAT) (4). Of these, the RCT and VAT are the most widely used physiological techniques for examining SCC function in clinical practice. In contrast to the caloric test, the RCT and VAT enable examination of the VOR at different frequencies ranging from 0.01 to 0.64 Hz and 2 to 6 Hz, respectively. Both tests provide important diagnosis information in terms of detecting overall function of the bilateral peripheral vestibular system. However, they are unlikely to provide accurate diagnosis in patients with unilateral vestibular hypofunction (5-7). Therefore, it is necessary to explore the mechanism underlying the angular motion sensed by three pairs of SCCs simultaneously to gain a better understanding of the clinical phenomena by which the SCCs interact in association with head movements.

Several models have been developed to explore the correlation between vestibular responses and/or neural responses and angular acceleration since the 1920s. For example, SCC fluid mechanics were first simulated by Lorente de Nó (8). In this model, an expression for transient endolymph flow in response to a step change in head velocity was derived. A classical torsion-pendulum model was developed by Steinhausen (9) to investigate the cupular volume displacement relative to the angular velocity of the head. Since then, several modified torsion-pendulum models and/or improved solution methods have been widely used to investigate the dynamic responses in SCCs to rotational stimulation (10-14). So far, both single-canal and three-canal models have been well developed to explore endolymph-cupula interaction during three-dimensional motions of the head. Obrist (15) concluded that the cupular displacement obtained from the torsion-pendulum models is roughly proportional to the head rotation velocity, as well as the nystagmus velocity, and thus, the volume displacement is a good means of expression for cupular dynamics.

In the past decades, the finite element (FE) method has become an alternative approach to investigate the biomechanical behaviors of SCCs, because this method better reflects the physiopathological characteristics and actual structure of the SCCs by providing biomechanical properties and geometrical parameters of the SCCs (16). Compared to torsion-pendulum models, in FE models, both detailed endolymphatic flow and cupular displacement responses to the head rotation are well developed and characterized (17, 18). However, evidence shows that the cupular displacement varies in both longitudinal and transverse directions (17-19). Such discrepancies were found in the torsion-pendulum models and FE models (i.e., although the direction of the displacements using FE models is the same as the displacements using torsion-pendulum models, both the magnitude and the phase of the displacements are not comparable).

As indicated above, although a series of responses to the head rotation have been simulated and measured, to our best knowledge, until now, there are no agreed-upon mechanical variables used in various models for detecting the angular motion and its mechanism in association with the function of three pairs of SCCs.

In the present study, an FE model of bilateral SCCs was developed. The complex fluid-structural interaction between the endolymph and cupulae was simulated and analyzed during sinusoidal head rotation by calculating the degree of cupular expansion and the cupular deflection. The mechanism of angular motion in association with the function of three pairs of SCCs was also explored.

### Materials and Methods

### Three-dimensional reconstruction of the bilateral vestibular labyrinth

The dimensions of a healthy adult inner ear membranous labyrinth were extracted from the data reported by Ifediba (20). Using these data, the pars superior of the membranous labyrinth in the right ear including three SCCs (the horizontal SCC [HC], the anterior SCC [AC], and the posterior SCC [PC], and the utricle was reconstructed using the FE software ANSYS (Ansys Inc., Canonsburg, PA, USA), as shown in Figure 1A. The pars superior in the left ear was also built using the symmetry of human bilateral ears along the sagittal plane (Figure 1A). In this model, the thickness of the membranous labyrinth was set to 30 µm (21), and three ampulla cupulae were assumed to be perpendicular to the longitudinal axis direction. The space surrounding the surface of three cupulae and the membranous labyrinth were filled with endolymph, and the effect of the macula utriculi was not considered. In the present study, the utriculo-endolymphatic valve (UEV) was assumed to stay closed and was incorporated into the surface of the membranous labyrinth due to the fact that it is usually considered to be closed to maintain the normal endolymph volume in the pars superior (22).

**Figure 1.**
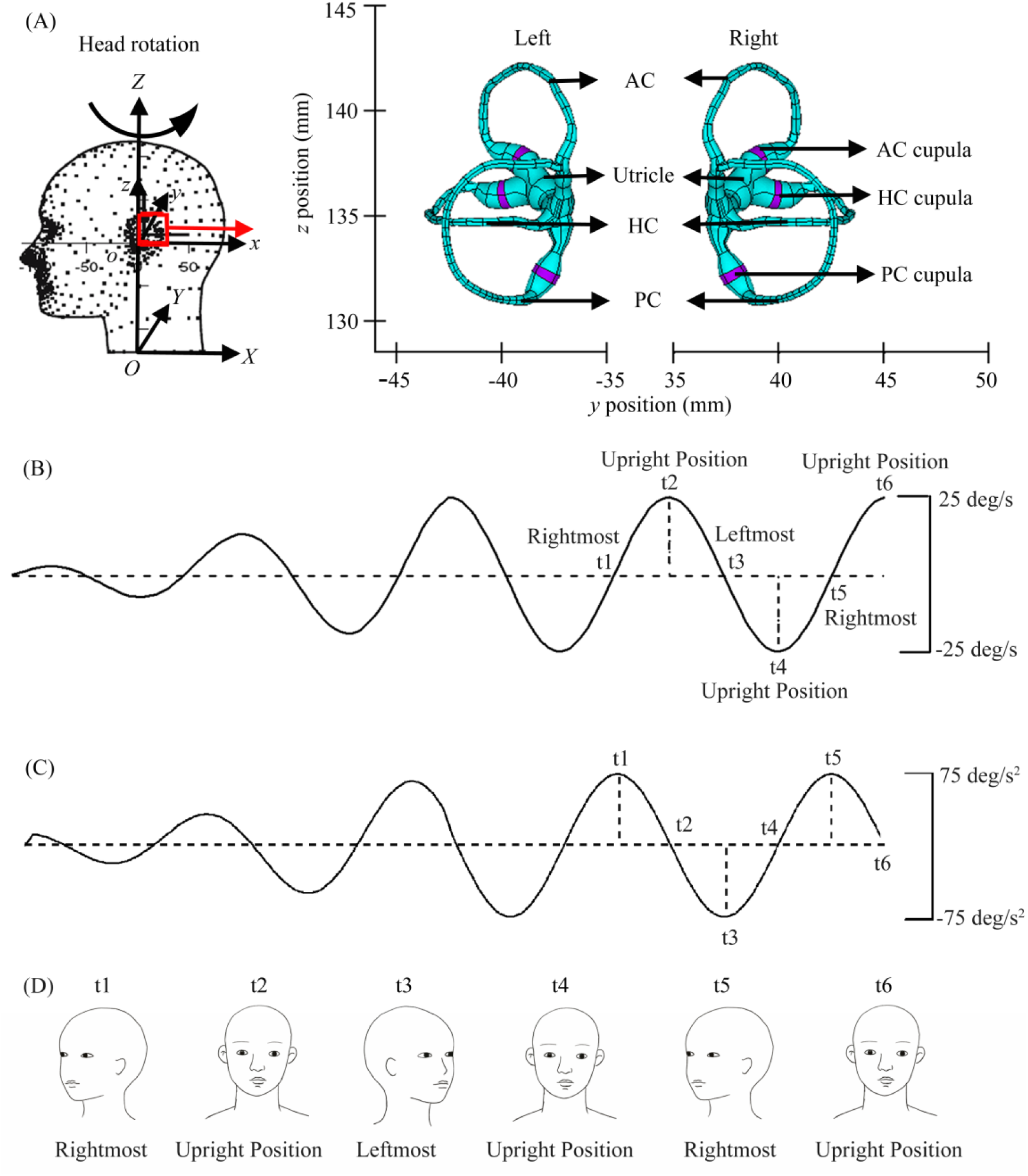
The computational model of angular motion in association with bilateral semicircular canal function. Figure 1(A) is the reconstructed geometry; HC, AC and PC are the horizontal SCC, the anterior SCC and the posterior SCC, respectively; Figure 1(B) is the horizontal sinusoidal head rotation around the Z-axis with an angular velocity amplitude of 25 deg/s but at different frequencies of 0.5 Hz, 1 Hz, 2 Hz, 4 Hz, and 6 Hz. At the beginning, the head is in the upright position, the head-fixed coordination (xyz) and the ground-fixed coordination (XYZ) are coincident with each other, with the X-axis toward the back of the head, the Y-axis toward the right side of the hand, and the Z-axis toward the top of the head; t1∼t6 are the different positions during head rotation, with the head passing the upright position at the t2, t4, and t6 positions, arriving at the rightmost position at the t1 and t5 positions, and arriving at the leftmost position at the t3 position. Figure 1(C) is the horizontal sinusoidal head rotation around Z-axis with an angular acceleration amplitude of 75 deg/s^2^ but at different frequencies of 0.5 Hz, 1 Hz, 2 Hz, 4 Hz, and 6 Hz. Various loading modes are used to explore the effect of angular velocity and angular acceleration on the frequency characteristics of cupular mechanical responses. Figure 1(D) is the sketch map of the head rotation corresponding to the positions of t1∼t6.

### Finite element model development

A sinusoidal horizontal head rotation was utilized to explore the mechanism of angular motion in association with three pairs of SCCs. The FE model of the SCCs was initially developed using the ADINA software (v8.7.1, ADINA R&D Inc.). The endolymph was simulated by using slightly compressible Newtonian fluid, which was divided using tetrahedral fluid elements. For the solid domain, the cupula was modeled using tetrahedral solid elements. However, the surface of the membranous labyrinth and the UEV were modeled using triangular shell elements, characterized as linearly elastic material undergoing large deformation and involving small strain. A strict grid sensitivity analysis was conducted and reported in our previous study (17). Finally, a spatial resolution of 100 µm was adopted to discretize the inner ear membranous labyrinth. The material properties used in this study were extracted from previous publications, as shown in Table 1, except the density of the membranous labyrinth, which was assumed to be the same as that for the cupula.

**Table 1.**
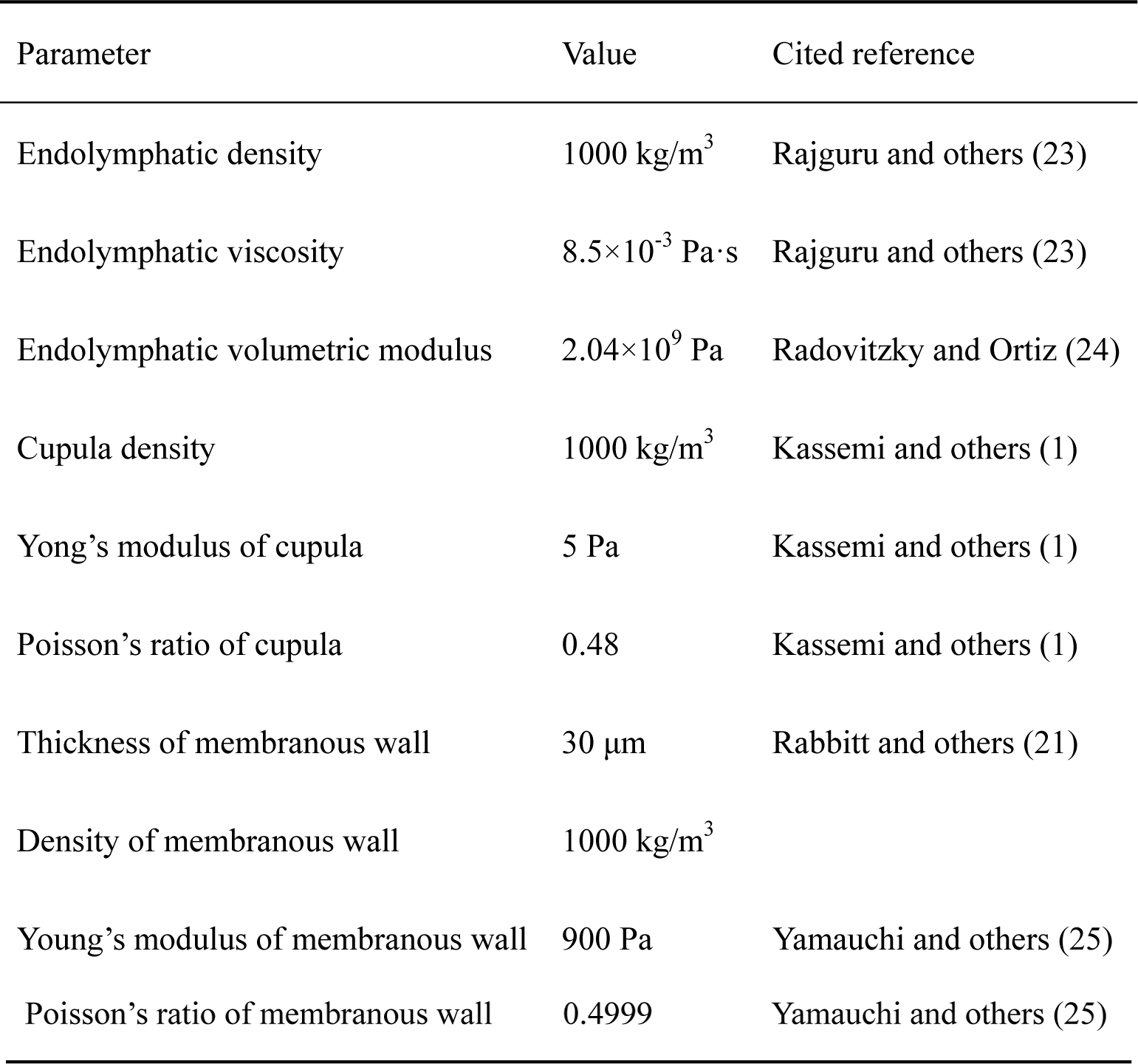
Physical properties of the SCC system used in the FE model

| Parameter | Value | Cited reference |
| --- | --- | --- |
| Endolymphatic density | 1000 kg/m<sup>3</sup> | Rajguru and others (23) |
| Endolymphatic viscosity | 8.5×10<sup>-3</sup> Pa·s | Rajguru and others (23) |
| Endolymphatic volumetric modulus | 2.04×10<sup>9</sup> Pa | Radovitzky and Ortiz (24) |
| Cupula density | 1000 kg/m<sup>3</sup> | Kassemi and others (1) |
| Yong's modulus of cupula | 5 Pa | Kassemi and others (1) |
| Poisson's ratio of cupula | 0.48 | Kassemi and others (1) |
| Thickness of membranous wall | 30 $\mu\text{m}$ | Rabbitt and others (21) |
| Density of membranous wall | 1000 kg/m<sup>3</sup> |  |
| Young's modulus of membranous wall | 900 Pa | Yamauchi and others (25) |
| Poisson's ratio of membranous wall | 0.4999 | Yamauchi and others (25) |

The interaction between the endolymph and all cupulae, as well as the surface of the membranous labyrinth and the UEV, were calculated using the following equations. For the fluid domain, the endolymph flow was calculated using the continuity and momentum equations, respectively:

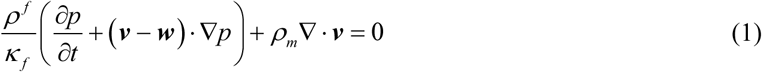

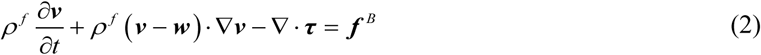

where *ρ* ^*f*^ is the endolymph reference density, *κ* ^*f*^ is the bulk modulus, *p* is the pressure, *t* is the time, ***v*** is the flow velocity vector, ***w*** is the mesh velocity vector, and ***f*** ^B^ is the body force per unit volume. Regarding the slightly compressible density *ρ*_*m*_ and the stress tensor ***τ***, these parameters were calculated using equations (3) and (4):

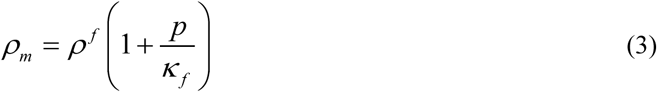

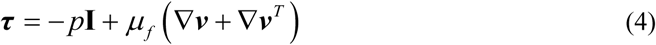

where **I** is the identity matrix and *μ*_*f*_ is the endolymph viscosity. For the solid domain, the motion of all solid structures was calculated using the Navier equation:

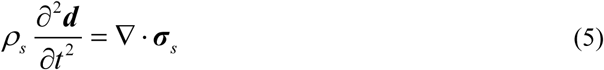

In equation (5), *ρ*_*s*_ is the density, and ***d*** is the displacement vector. The stress tensor ***σ***_*s*_ was defined as follows:

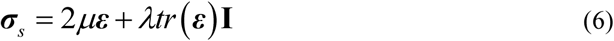

where ***ε*** is the strain tensor, and *μ* and *λ* are Lamé coefficients, which are related to Young’s modulus of elastic *E* and Poisson’s ratio *ν*, according to the following equations:

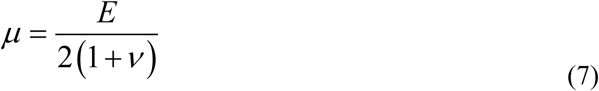

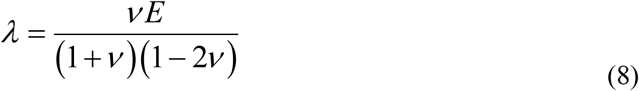

At the beginning, the head was assumed to be static and in the upright position. Then, as shown in Figure 1B, a progressive sinusoidal wave was applied in the first two cycles to avoid problems with discontinuity and improve numerical reliability and convergence, followed by undergoing a stationary sinusoidal wave with a maximum angular velocity of 25 deg/s. The frequency characteristics responding to the rotations in the SCCs were analyzed at frequencies of 0.5 Hz, 1 Hz, 2 Hz, 4 Hz, and 6 Hz, respectively, which are close to the frequency range of head movement in daily life.

Furthermore, to compare the effect of angular velocity to that of angular acceleration on the cupular mechanical responses, the same series of sinusoidal waves were used, but with the maximum angular acceleration of 75 deg/s^2^, as shown in Figure 1C. By using the fine filaments, the surface of the membranous labyrinth was considered to keep pace with the bony labyrinth as well as the head. As a result, intricate interactions between the endolymph and all cupulae occurred due to inertia with wetted coupling interfaces satisfying a balance of traction forces and continuity of velocities and displacements.

The fluid and solid equations were eventually coupled to calculate on the basis of the direct solution procedures using the arbitrary Lagrange-Euler (ALE) approach. The time steps at each rotation period were consistently set to 50. At each time step, the convergence tolerance for the velocity and displacement norms was 0.001, and for the fluid-solid surface norm, it was 0.01. The reference value was automatically chosen by the program. As shown in Figure 1A, the relative cupular displacement in the head-fixed coordinate system (xyz), defined to align with the ground-fixed system (XYZ) when the subject is in the upright position prior to movement of the head, was rigorously examined. Both the endolymphatic flow and the relative cupular node displacement were closely examined in our previous report in terms of their verification and reliability (17). In the present study, the angular motion in association with the three SCCs was characterized by the cupular deflection *d*_*c*_ and the degree of cupular expansion Δ and can be defined by equations (9) and (10), respectively:

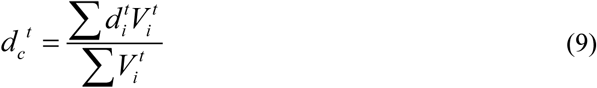

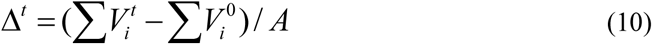

Where 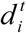 and 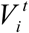 denote the relative centroid displacement and the volume of the *i*^th^ tetrahedral element at moment *t*, respectively; Δ*t* is the degree of cupular expansion at moment *t*; and 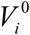 is the volume of the *i*^th^ tetrahedral element at the beginning of head rotation. *A* is the area of the middle cross section for the HC cupula, the AC cupula, and the PC cupula, respectively. Both the degree of cupular expansion and the cupular volumetric deflection were calculated using MATLAB software.

## Results

### The responses of the degree of cupular expansion to sinusoidal head rotation

When the head made any rotation, three pairs of SCCs in both left and right ears were simultaneously stimulated. Figure 2 shows the responses of the degree of expansion for three pairs of bilateral symmetrical cupulae to sinusoidal rotation of 25 deg/s at the frequency of 2 Hz. All expansion curves of the SCCs positively or negatively correlated with the head rotation. The responses of the left HC cupula and the right HC cupula almost coincided (i.e., they expanded or compressed simultaneously), as did the bilateral AC cupula and the bilateral PC cupula. Moreover, when the head rotated to the right (i.e., at the positions of t1∼t2 and t4∼t5, shown in Figures 1B and 1D), bilateral HC and AC cupulae expanded, both PC cupulae compressed, and vice versa. In addition, all expansion curves were completely fitted by sinusoidal functions, as in the following formala:

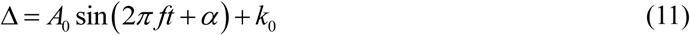

**Figure 2.**
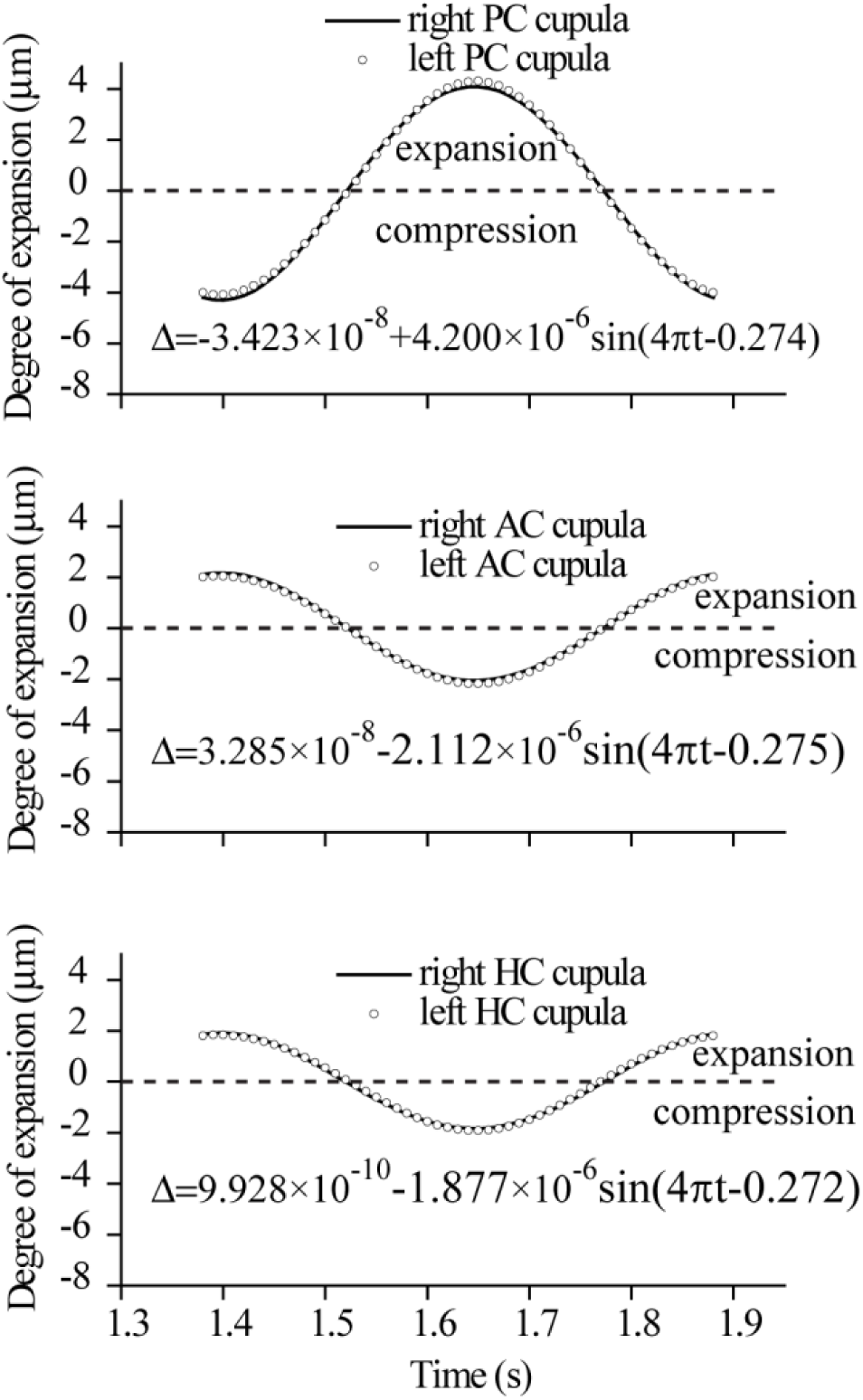
The responses of the degree of cupular expansion to sinusoidal rotation of 25 deg/s at the frequency of 2 Hz at the positions of t1∼t5, shown in Figure 1. It is positive when the cupula dilates, and vice versa. The responses for bilateral symmetrical cupulae almost coincide, and the degree of expansion for the HC, AC, and PC cupulae can be fitted by a sinusoidal function with the same frequency of head rotation, respectively.

where *A*_*0*_ is the amplitude; *f* is the frequency, which equals the stimulating frequency, *α* is the initial phase; and *k*_0_ is the intercept.

In this study, the expansion amplitude for the PC cupula was approximately 4 *µ*m, which was about two times greater than those for the HC and AC cupulae. In addition, their phases uniformly led the angular displacement of approximately 15.5 degrees, but they were in the reverse phases to those for both HC and AC cupulae.

The frequency characteristics responding to the rotations in three right cupular expansions were further analyzed, as shown in Figure 3. When the head made a series of sinusoidal rotations with the same angular velocity amplitude of 25 deg/s, but at different frequencies of 0.5 Hz, 1 Hz, 2 Hz, 4 Hz, and 6 Hz, respectively, the expansion amplitude of the three cupulae increased linearly with the increase of in the frequency. However, there was a plateau at the frequency band between 2 Hz and 4 Hz (Figure 3A). In addition, the amplitude of the PC cupula was the strongest at all frequencies, which were approximately two times greater than those of both the HC and AC cupulae. On the other hand, however, their phase only decreased linearly with the increase of frequency (Figure 3B). The phase of the HC cupular expansion was equal to those of the AC cupula, but both of them differed by 180 degrees from those of the PC cupula.

**Figure 3.**
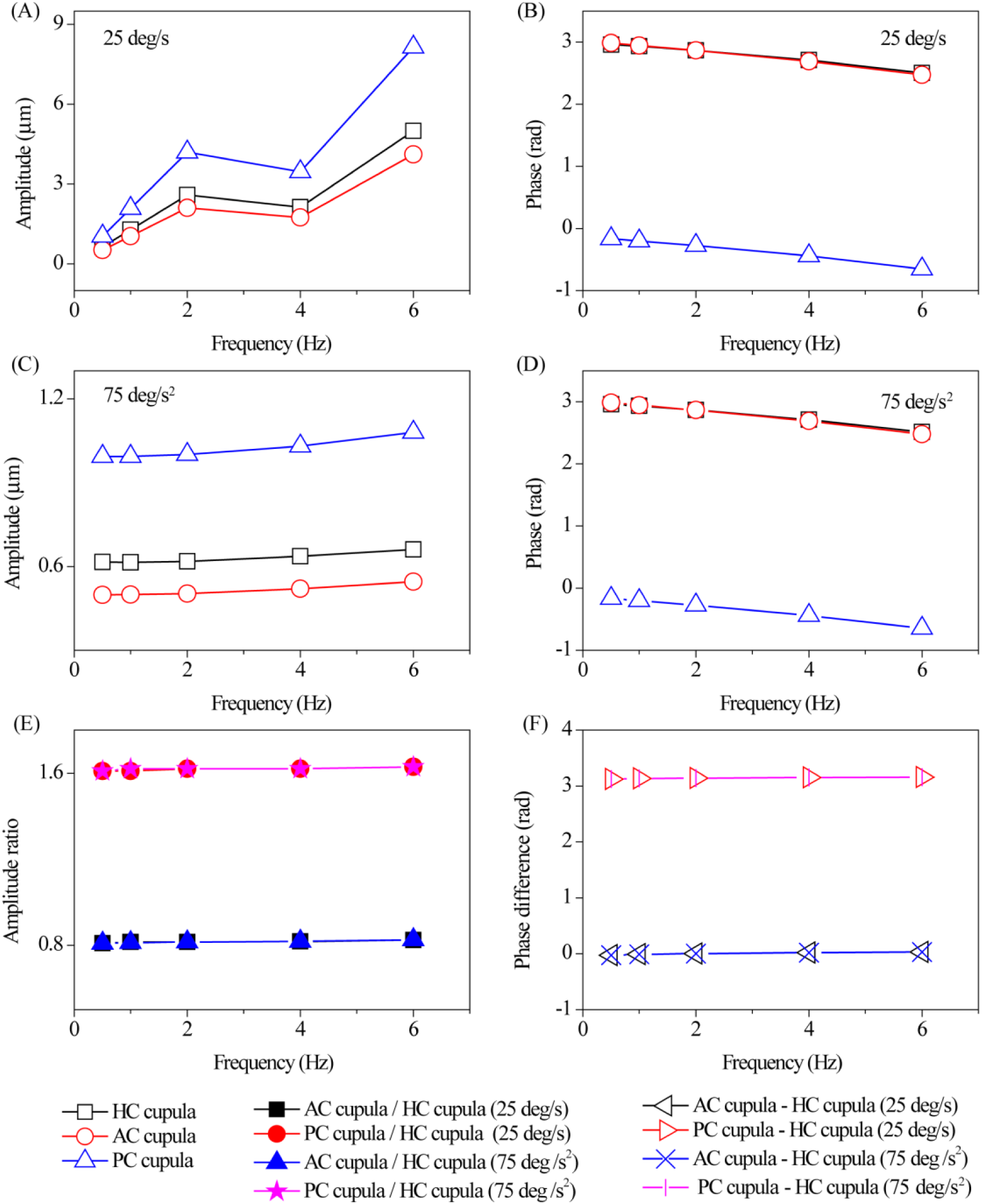
The frequency characteristics of the degree of cupular expansion. Figures 3(A) and 3(B) are the amplitude-frequency characteristic and the phase-frequency characteristic during head horizontal sinusoidal rotation with the same velocity amplitude of 25 deg/s and at different frequencies of 0.5 Hz, 1 Hz, 2 Hz, 4 Hz, and 6 Hz, respectively. Figures 3(C) and 3(D) are the amplitude-frequency characteristic and the phase-frequency characteristic during head horizontal sinusoidal rotation with the same acceleration amplitude of 75 deg/s^2^ and at different frequencies of 0.5 Hz, 1 Hz, 2 Hz, 4 Hz, and 6 Hz, respectively. Finally, Figures 3(E) and Figure 3(F) are the amplitude ratio characteristic and phase difference characteristic of the above two loading conditions, respectively.

Similarly, when the head made a series of sinusoidal rotations with the same angular acceleration amplitude of 75 deg/s^2^, but at different frequency of 0.5 Hz, 1 Hz, 2 Hz, 4 Hz, and 6 Hz, respectively, the expansion amplitude of the degree of the three cupulae also increased with the inrease of frequency, with the biggest amplitude occurring in the PC cupula and the smallest in the AC cupula (Figure 3C). In addition, the phase characteristics were found to be the same as when an angular acceleration amplitude of 25 deg/s^2^ was used (Figure 3D).

When the amplitude-frequency and the phase-frequency characteristics of the above two conditions in association with the head rotation were compared, the amplitude ratios of the HC cupular expansion, AC cupular expansion, and PC cupular expansion were constant at 1:0.82:1.62 for any frequency sinusoidal rotation stimulation (Figure 3E). Regarding the phase characteristics, the phase differences between the HC cupular expansion and AC cupular expansion were constant at 0, whereas those btween the HC cupular expansion and PC cupular expansion were constant at 180 degrees. Figure 3F further demonstrates that the phase for the HC cupula and AC cupula was the same, but they were out of phase to those of the PC cupula at any frequency of sinusoidal rotation stimulation.

### The responses of cupular volumetric deflection to sinusoidal head rotation

As shown in Figure 4, although they still positively or negatively correlated with the head rotation, the deflection responses of the left HC cupula and right HC cupula were opposite in phase (i.e., when the right HC cupula deflected to the utricle side, the left HC cupula almost equally deflected to the canal side). The same responses were found in the other two pairs of cupulae. In addition, all deflection curves were fitted well to sinusoidal functions, with the amplitude of each pair’s cupular deflection equal, according to the following formula:

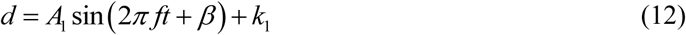

where *A*_*1*_ is the amplitude, *β* is the initial phase, and *k*_1_ is the intercept. Specifically, when the head rotated to the right (i.e., at the positions of t1∼t2 and t4∼t5, shown in Figure 1B), the right HC cupula, the right PC cupula, and the left AC cupula produced exciting stimulation, but the other cupulae produced inhabitation stimulation, and vice versa. The defection amplitude for both PC cupulae was approximately 2 µm, which was also about two times greater than those for both the HC and AC cupulae. In the phase analysis, the phases found in the SCCs were not exactly identical to each other (i.e., the cupular deflection of the left HC cupula led to angular displacement of about 10.6 degrees, and those of the right AC cupula and the left PC cupula lagged the angular displacement by 8.3 degrees and 17.6 degrees, respectively). Moreover, the phases differed by 180 degrees from their symmetrical cupulae.

**Figure 4.**
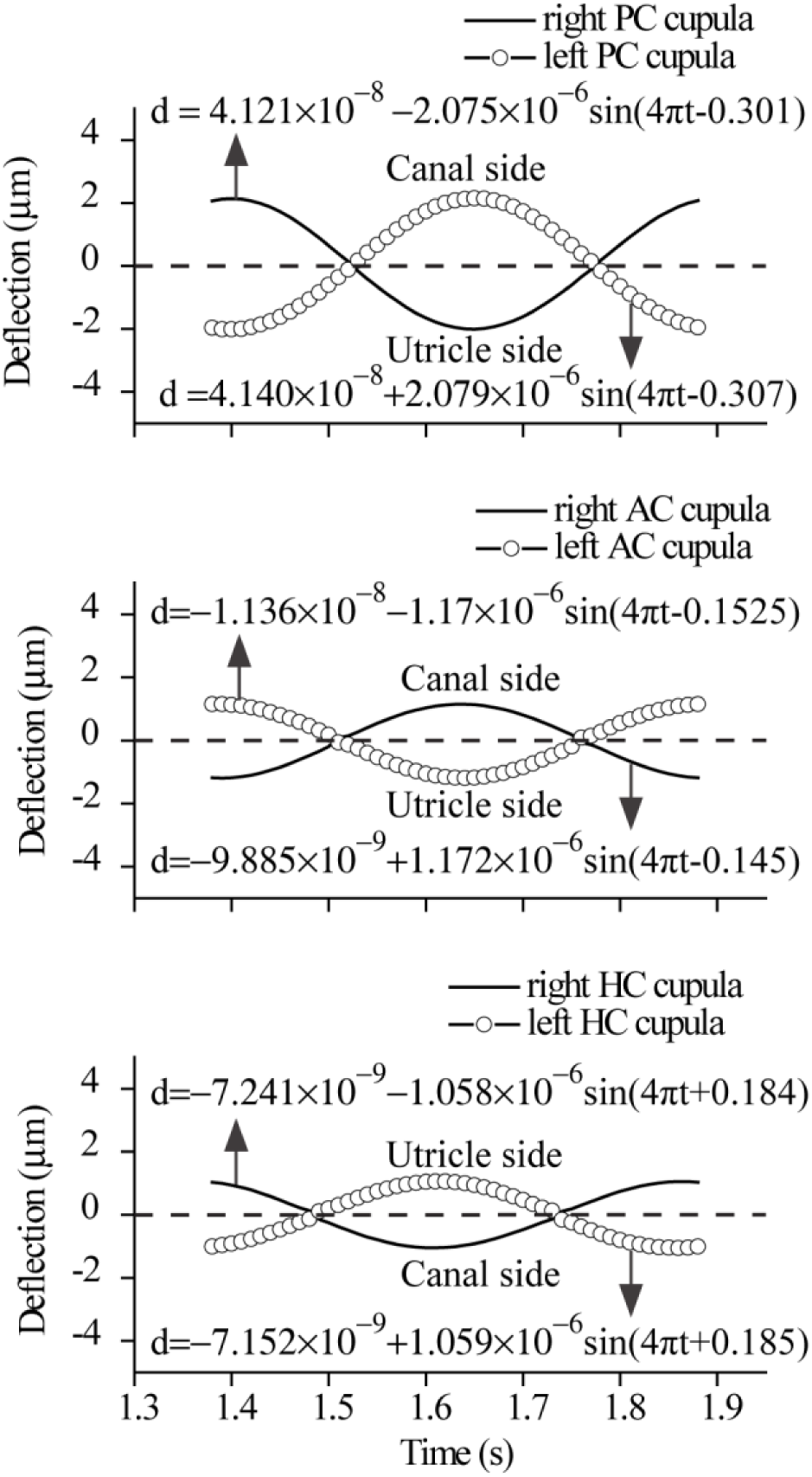
The responses of the cupular deflection to sinusoidal rotation of 25 deg/s at the frequency of 2 Hz at the positions of t1∼t5, shown in Figure 1. The cupular deflection responses is positive when the deflection makes the sensory cilium produce excited signals (i.e. the HC cupula deflects to the utricle side, whereas the other cupulae deflect to the canal side, and vice versa). Specifically, when the right cupula is excited, the left corresponding one will be inhibited.

The frequency characteristics responding to the rotations in the three right cupular deflections were further analyzed (Figure 5). When the head made a series of sinusoidal rotations with the same angular velocity amplitude of 25 deg/s, but at different frequencies of 0.5 Hz, 1 Hz, 2 Hz, 4 Hz, and 6 Hz, respectively, the amplitude-frequency characteristics of cupular deflection were extremely similar to those of cupular expansion (i.e., they also increased linearly with the increase of frequency). Similarly, there was a plateau at the frequency band between 2 Hz and 4 Hz, and the amplitude of the PC cupula was the greatest at all frequencies, which were approximately two times greater than those of the other two cupulae at the high frequency band between 2 and 6 Hz (Figure 5A). For the phase analysis, the phase generally decreased with the increase of frequency but showed little characteristic tendency. There were larger phase differences at the frequencies of 0.5 to 2 Hz than at the frequencies of 2 to 6 Hz. The HC and PC cupular deflections were in phase, but they were out of phase to the AC cupular deflection.

**Figure 5.**
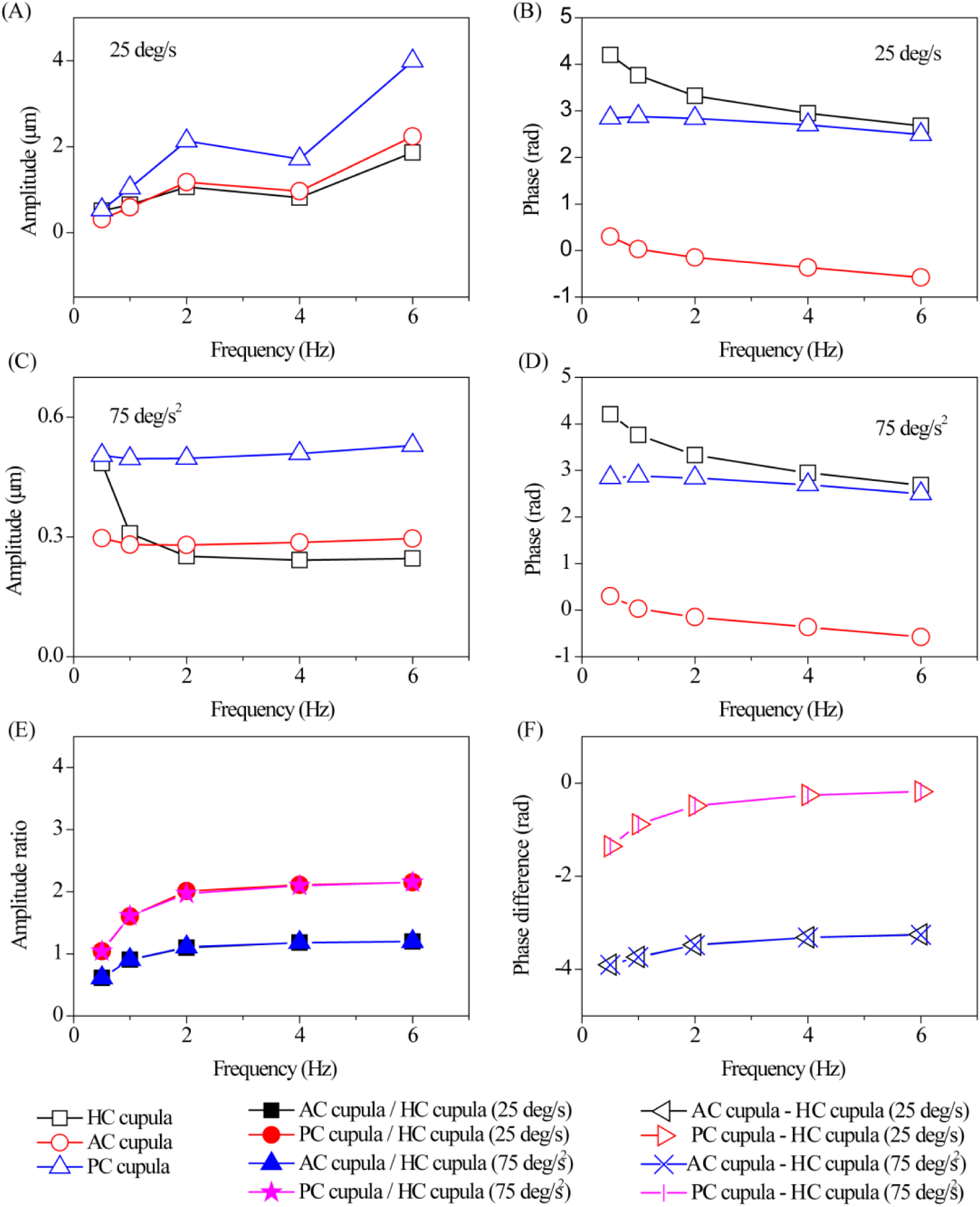
The frequency characteristics of the cupular deflection. Figures 5(A) and 5(B) are the amplitude-frequency characteristic and the phase-frequency characteristic during head horizontal sinusoidal rotation with the same velocity amplitude of 25 deg/s and at different frequencies of 0.5 Hz, 1 Hz, 2 Hz, 4 Hz, and 6 Hz, respectively. Figures 5(C) and 5(D) are the amplitude-frequency characteristic and the phase-frequency characteristic during head horizontal sinusoidal rotation with the same acceleration amplitude of 75 deg/s^2^ and at different frequencies of 0.5 Hz, 1 Hz, 2 Hz, 4 Hz, and 6 Hz, respectively. Finally, Figures 5(E) and 5(F) are the amplitude ratio characteristics and phase difference characteristics of the above two loading conditions in association with the head rotation, respectively.

Furthermore, when the head made a series of sinusoidal rotations with the same angular acceleration of 75 deg/s^2^, but at different frequencies of 0.5 Hz, 1 Hz, 2 Hz, 4 Hz, and 6 Hz, respectively, the amplitude of the three cupular deflections first decreased with the increase of frequency at the frequency band between 0.5 and 2 Hz, and then increased with the increase of frequency at frequency band between 2 and 6 Hz. The largest amplitude occurred in the PC cupula and the smallest one in the AC cupula (Figure 5C). In addition, the phase characteristics were found to be the same as when an angular acceleration amplitude of 25 deg/s^2^ (Figure 5D).

When the amplitude-frequency and phase-frequency characteristics of the above two conditions in association with the head rotation were compared, the amplitude ratios among the HC cupular deflection, AC cupular deflection, and PC cupular deflection increased nonlinearly with the increase of frequency, and it tended to a constant of 1:1.2:2.15 for sinusoidal rotation stimulation frequencies larger than 2 Hz (see Figure 5E). Regarding the phase characteristics, the phase differences between AC cupular deflection and HC cupular deflection and those between PC cupular deflection and HC cupular deflection also nonlinearly increased with the increase of frequency, and they tended to constants of −187 degrees and −11 degrees, respectively, for sinusoidal rotation stimulation frequencies larger than 2 Hz (see Figure 5F).

## Discussion

In the present study, we fully used the advantages of both the lumped torsion pendulum model and the FE model, and consequently, novel biomechanical analyses were performed using the FE model by calculating the degree of expansion and the deflection of the whole cupula. The advantage of the present model is that it not only ensures that the physical and geometrical parameters of SCCs are as accurate as possible to their physiological condition, but it also measures the cupular responses in association with head rotation as a unique solution to the forced vibration. Therefore, the quantitatve relationship between the head rotation and the mechanical responses of SCCs is established. Essentially, endolymphatic flow would apply two types of pressure on the two sides of the cupula (i.e., the expansion pressure Δ*p* and the pressure difference *dp*):

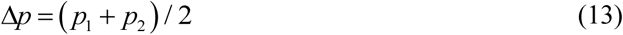

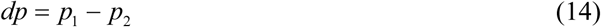

Where *p*_1_ and *p*_2_ are the pressure on the utricle side and the canal side of the cupula, respectively. The cupula generates the responses of the degree of expansion due to the expansion pressure, and it generates deflection responses due to the pressure difference. As a result, it is important to analyze the effect of both the degree of cupular expansion and cupular deflection on the cupular mechano-electrical transduction process at the same time.

In previous studies, cupular deflection was usually considered a mechano-electrical transduction variable. For example, according to Ewald’s II law, when the head horizontally rotates to the right (at positions of t1∼t2 and t4∼t5, shown in Figures 1B and 1D), the endolymph in the right ear would generate ampullopetal flow due to inertia, and the right HC cupula would deflet to the utricle side to generate stimulating signals. In contrast, the left HC cupula would deflect to the canal side to generate inhibited signals, and vice versa. The present results are completely in keeping with Ewald’s II law (as shown in Figure 4).

Theoretically, when the cupula is expanded, the distance between any two points along the longitudinal direction, as well as the distance between the stereocilia and kinocilia, would be elongated, and consequently, it generates stimulating signals for the mechano-electrical transduction process, and vice versa. Therefore, the degree of cupular expansion plays an important role in the cupular mechano-electrical transduction process, although there has been little research until now.

As found in this study, the maximum value of the degree of cupular expansion is approximately two times greater than that of the corresponding cupular deflection (shown in Figurs 2 and 4). Therefore, the effect of cupular expansion on the mechano-electrical transduction should be further investigated in terms of the association with some clinical vestibular examinations, such as both the RCT and VAT, particularly regarding the connection to the eye movements via VOR. The sketch map of the VOR pathway is shown in Figure 6.

**Figure 6.**
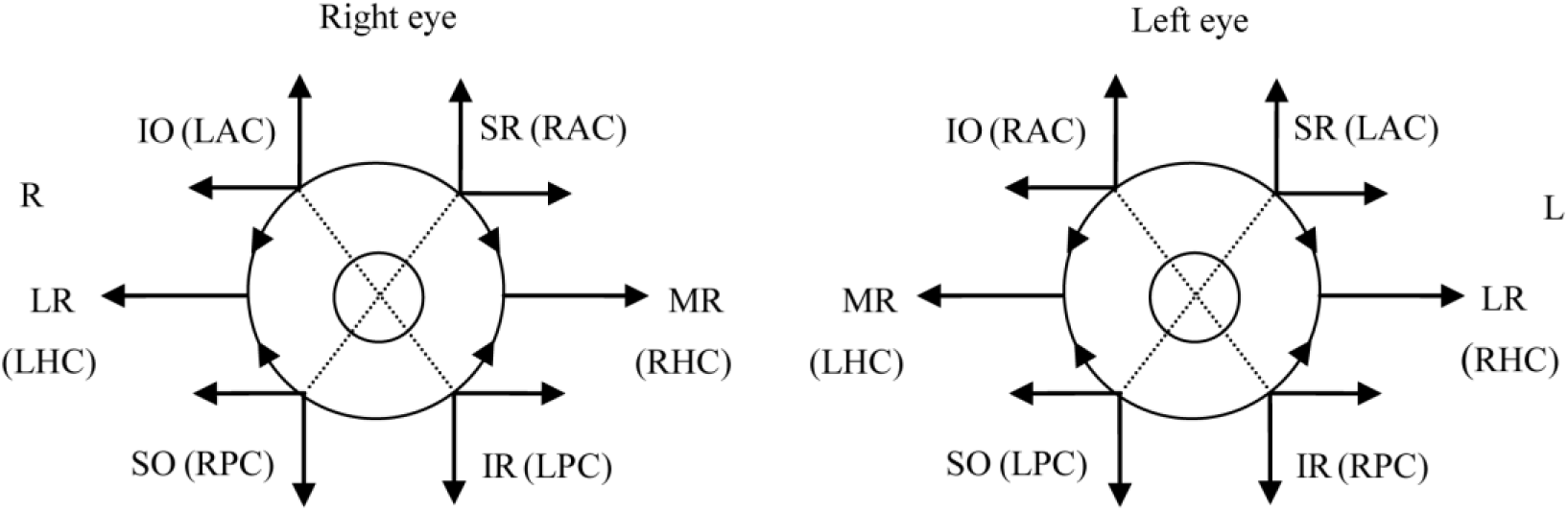
The relationship among three pairs of SCCs, extraocular muscles, and eye movements via the VOR pathway. LR: the lateral rectus, MR: the medial rectus, SR: the superior rectus, IR: the inferior rectus, SO: the superior oblique, IO: the inferior oblique, RHC: the right horizontal semicircular canal, LHC: the left horizontal semicircular canal, RAC: the right anterior semicircular canal, LAC: the left anterior semicircular canal, RPC: the right posterior semicircular canal, LPC: the left posterior semicircular canal.

According to the results obtained in this study, when the head rotates to the right, both HC cupulae and both AC cupulae, respectively, expand equivalently, and both PC cupulae are compressed equivalently. As a result, according to the VOR pathway, both the evoked horizontal eye movement and the evoked rotational eye movement are always in a state of equilibrium (see Figure 6). For the evoked vertical eye movement, it is assumed that the downward turning resulted from the inward turning and the outward turning of both AC cupulae can balance out the upward turning, and so do both PC cupulae. When the head rotates to the left, the eye movement in all directions can also reach a balance. However, if the balance is destroyed by any abnormalities of inner anatomical structure or function, the balance of the degree of cupular expansion is also damaged, and the final relative eye movement will be the sum of the motion evoked by both the degree of cupular expansion and cupular deflection.

Furthermore, the contribution of the cupular deflection was also analyzed. Based on the results found in this study, when the head rotates to the right, the right HC cupula, the right PC cupula, and the left AC cupula are stimulated unequally, and the other three cupulae are inhibited. According to the VOR pathway, the left turning driven by the medial rectus (MR) and the right HC cupula is partially counteracted by the right turning driven by the inferior oblique (IO) and the superior oblique (SO); upward turning driven by the IO is also partially counteracted by downward turning driven by the SO, as well as the inward turning and outward turning. Therefore, it can be predicted that when the rotation angle of the left turning for the eyeball reaches a certain level, the movement will stop with the function of both the mechanical force of six ocular muscles and the force produced from the VOR. It might constitute the slow phase of the eye movement, and then the fast phase would occur due to central regulatory. It is worth mentioning that the up or down vertical turning and the outward or inward rotation of the eyeball are actually very small in the normal case when the head undergoes horizontal rotation. However, when the head rotates to the left, the direction of the eye movement is the exact opposite.

In this study, both the degree of cupular expansion and cupular deflection can be expressed as the solution of forced oscillation during head sinusoidal rotation, and both the amplitude and phase can be further expressed as the function of the mass, the stiffness and the damping of the vibration system. Specifically, when the head is subjected to any horizontal sinusoidal head rotation, the following equation applies:

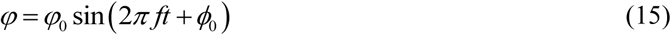

where *φ*_*0*_ and *ϕ*_*0*_ are the amplitude and the initial phase of the head rotation, respectively. Both the degree of the cupular expansion and the cupular deflection can be expressed by formulae (11) and (12), respectively. According to the basic principle of the forced vibration, the amplitude of responses *A*_*0*_ and *A*_*1*_ can be uniformly represented by A as follows:

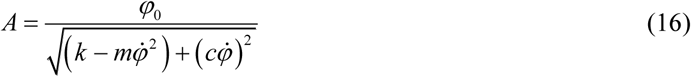

and the initial phase can be expressed by the following equation:

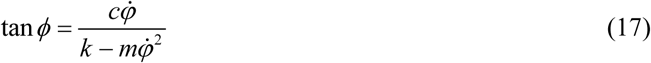

where *k, m* and *c* are the stiffness, the mass and the damping of the vibration system (i.e. the SCCs system), respectively.

In fact, the frequency characteristic of the SCC system with respect to sinusoidal head rotation was first discovered in the 1970s by introducing the torsion-pendulum model (26), and the responses are quantitatively represented by the amplitude, the phase, and the time constants. These eigenvalues are usually used for clinical diagnosis. The forced vibration pattern further demonstrates that our model is reliable. Even so, the FE model presented in this study could not be directly compared to the torsion-pendulum model presented by Fernandez or other models due to the fact that the SCCs are applied with different body positions, rotation axes, and initial phases. Thus, the function of SCCs can be affected by the mass, the stiffness, and the damping of the SCC system, and then they feed back to the amplitude, the phase, and the symmetry of the eye movements and clinical diagnosis.

The effect of the physical properties on cupular responses can be roughly divided into several aspects according to the forced vibration theory. If something causes the stiffness of the SCC system larger than the normal range, it will lead to both the amplitude and the phase being smaller than the normal range (see equations 16 and 17), as well as the eye movement, and vice versa. Similarly, if something causes the mass of the SCC system to be larger than the normal range, both the amplitude and the phase of cupular responses are also smaller than the normal range, as well as the eye movement, and vice versa. If something causes the damping of the SCC system to be larger than the normal range, it will result in the amplitude of the cupular responses to be larger than the normal range, but the phase will become smaller, and vice versa. If the difference is in the normal medical reference range, the vestibular rotational tests cannot indicate any dysfunction of SCCs. Overall, any vestibular disorders might simultaneously affect the above two or three parameters, and the role of the type and severity of the vestibular disease in quantitatively affecting the SCCs’ function needs to be further explored.

In addition, the frequency characteristics of the SCC system in response to sinusoidal head rotation can be divided into the frequency characteristics of the degree of cupular expansion and cupular deflection. In general, the frequency characteristic of the degree of cupular expansion is more significant, and not only do the phases of three cupular expansion remain the same or opposite, but the amplitude ratios among them remain constant, and the amplitude ratios of the HC, the AC, and the PC cupular expansion is 1:0.82:1.62. This implies that the degree of cupular expansion is affected by both angular acceleration and angular velocity simultaneously, but their quantitative relationship is probably only related to the spatial orientation among the three SCCs. From the view of mechanics, the vestibular system can be considered a linearly undamped forced vibration system, and the degree of cupular expansion might only be related to the mass and the rigidity of the three cupulae and the endolymph.

However, regarding the frquency characteristics of cupular deflection, neither the amplitude feature nor the phase feature is significant, especially at the low freqeucny band of 0.5 to 2 Hz but tends to be constant at a higher frequency band between 2 and 6 Hz. The results presented in this study agree with the prediction of Su Haijun (27). It can be deduced that the amplitude of the responses among the three SCCs is not only related to their spatial orientation, but also might be affected by the other physical properties of the system, such as the mass, the rigidity, or the damping.

For future studies, it seems feasible to further build numerical models by modifying the mass, the rigidity, and the damping characteristics of the SCC system under different abnormal conditions to explore the effects of cupular expansion and deflection on the corresponding nystagmus in association with these pathological vestibular disorders. This would provide useful information contributing to a better understanding of the biomechanical nature underlying vestibular dysfunction.

## Conclusion

An FE model of healthy human vestibular SCCs in association with head sinusoidal rotation has been established. The mechanical responses were expressed as the degree of cupular expansion and cupular deflection. By using this model, the vestibular responses were analyzed, showing quantitative correlation with the eye movement due to the VOR pathway. Under normal conditions, the inverse eye movement is mainly caused by cupular deflection during head rotation, whereas little is affected by the degree of cupular expansion. In contrast, under abnormal conditions, when the balance of bilateral vestibular mechanical stimulation is destroyed, it leads to abnormal nystagmus due to the above two mechanical responses. Therefore, both cupular deflection and expansion should be considered important mechanical variables for induced neural signals. Numerical models can be further built by modifying the mass, stiffness, and damping of the SCC system, and consequently, they provide more theoretical diagnostic information about the nature and degree of vestibular dysfunction.

## Authors contributions

SS contributed largely to conduct execution of the numerical simulation, data analysis and manuscript writing. FZ provided suggestion on the study, and writing up the manuscript. ZYC, QYZ, SY, TTC and PM contributed to study design, data analysis and commented on result interpretation and discussion.

## Acknowledgements

The authors gratefully acknowledge support of this work from the National Natural Science Foundation of China (No. 31500765, 11572079, 81530030), the Foundation of Taishan Scholar (tshw20110515), and the 2019 Braille and Sign Language Project of China Disabled Persons’ Federations (CLS2019-02).

## References

1. Kassemi, M., D. Deserranno, and J. G. Oas. 2005. Fluid–structural interactions in the inner ear. Computers & Structures 83(2-3):181–189.

2. Burnette, E., E. G. Piker, and D. Frank-Ito. 2018. Reevaluating Order Effects in the Binaural Bithermal Caloric Test. American Journal of Audiology 27(1):104–109.

3. Morita, M., T. Imai, S. Kazunori, N. Takeda, I. Koizuka, A. Uno, T. Kitahara, and T. Kubo. 2003. A new rotational test for vertical semicircular canal function. Auris Nasus Larynx 30(3):233–237.

4. Hsieh, L.-C., T.-M. Lin, Y.-M. Chang, T. B. J. Kuo, and G.-S. Lee. 2015. Clinical applications of correlational vestibular autorotation test. Acta Oto-Laryngologica 135(6):549–556.

5. Baloh, R. W., K. Hess, V. Honrubia, and R. D. Yee. 1983. Low and High Frequency Sinusoidal Rotational Testing in Patients with Peripheral Vestibular Lesions. Acta Otolaryngol Suppl 96(sup406):189–193.

6. Baloh, R. W., S. M. Sakala, R. D. Yee, L. Langhofer, and V. Honrubia. 1984. Quantitative vestibular testing. Otolaryngology--head and neck surgery: official journal of American Academy of Otolaryngology-Head and Neck Surgery 92(2):145–150.

7. Hess, K., R. W. Baloh, V. Honrubia, and R. D. Yee. 2010. Rotational testing in patients with bilateral peripheral vestibular disease. Laryngoscope 95(1):85–88.

8. R, L. d. N. 1927. Contribution al estudio matematico del organo del equilibrio. Trabajo publicado en la 7:202–206.

9. Steinhausen, W. 1933. Über die Beobachtung der Cupula in den Bogengangsampullen des Labyrinths des lebenden Hechts. Pflügers Archiv Für Die Gesamte Physiologie Des Menschen Und Der Tiere 232(1):500–512.

10. Van Buskirk, W. C., R. G. Watts, and Y. K. Liu. 1976. The fluid mechanics of the semi-circular canals. Journal of Fluid Mechanics 78(1):87–98.

11. Oman, C. M., E. N. Marcus, and I. S. Curthoys. 1987. The influence of semicircular canal morphology on endolymph flow dynamics. An anatomically descriptive mathematical model. Acta Oto-Laryngologica 103(1-2):1–13.

12. Damiano, E. R., and R. D. Rabbitt. 1996. A singular perturbation model of fluid dynamics in the vestibular semicircular canal and ampulla. Journal of Fluid Mechanics 307(307):333–372.

13. Damiano, E. R. 1999. A poroelastic continuum model of the cupula partition and the response dynamics of the vestibular semicircular canal. J Biomech Eng 121(5):449–461.

14. Rabbitt, R. D. 1999. Directional coding of three-dimensional movements by the vestibular semicircular canals. Biological Cybernetics 80(6):417–431.

15. Obrist, D. 2011. Fluid Mechanics of the Inner Ear. Habilitation treatise in fluid mechanics at the department of mechanical and process engineering of ETH Zurich.

16. Kondrachuk, A. V., A. A. Shipov, T. G. Astakhova, and R. D. Boyle. 2011. Current trends in mathematical simulation of the function of semicircular canals. Human Physiology 37(7):802–809.

17. Shen, S., X. Sun, S. Yu, Y. Liu, Y. Su, W. Zhao, and W. Liu. 2016. Numerical simulation of the role of the utriculo-endolymphatic valve in the rotation-sensing capabilities of semicircular canals. Journal of Biomechanics 49(9):1532–1539.

18. Selva, P., J. Morlier, and Y. Gourinat. 2010. Toward a three-dimensional finite-element model of the human inner ear angular accelerometers sensors. International Journal for Computational Vision & Biomechanics.

19. Mclaren, J. W., and D. E. Hillman. 1979. Displacement of the semicircular canal cupula during sinusoidal rotation. Neuroscience 4(12):2001–2008.

20. Ifediba, M. A., S. M. Rajguru, T. E. Hullar, and R. D. Rabbitt. 2007. The role of 3-canal biomechanics in angular motion transduction by the human vestibular labyrinth. Annals of Biomedical Engineering 35(7):1247–1263.

21. Rabbitt, R. D., E. R. Damiano, and J. W. Grant. 2004. Biomechanics of the Semicircular Canals and Otolith Organs.

22. Hf, S., and B. AA. 1975. The utriculo-endolymphatic valve: its functional significance. The Jounal of Laryngology & Otology 89(10):985–996.

23. Rajguru, S. M., M. A. Ifediba, and R. D. Rabbitt. 2004. Three-Dimensional Biomechanical Model of Benign Paroxysmal Positional Vertigo. Annals of Biomedical Engineering 32(6):831–846.

24. Radovitzky, R., and M. Ortiz. 2015. Lagrangian finite element analysis of Newtonian fluid flows. International Journal for Numerical Methods in Engineering 43(4):607–619.

25. Yamauchi, A., R. D. Rabbitt, R. Boyle, and S. M. Highstein. 2002. Relationship between Inner-Ear Fluid Pressure and Semicircular Canal Afferent Nerve Discharge. Journal of the Association for Research in Otolaryngology 3(1):26–44.

26. Fernandez, C., and J. M. Goldberg. 1971. Physiology of peripheral neurons innervating semicircular canals of the squirrel monkey. II. Response to sinusoidal stimulation and dynamics of peripheral vestibular system. Journal of Neurophysiology 34(4):661–675.

27. Haijun, S., Y. Chunyu, W. Zhenguang, and X. Mingyu. 2005. Generalized fractional dynamic model of semicircular canal. Journal of Shandong University 40(1):37–41.

